# The fine-scale landscape of immunity and parasitism in a wild ungulate population

**DOI:** 10.1101/483073

**Authors:** Gregory F. Albery, Daniel J. Becker, Fiona Kenyon, Daniel H. Nussey, Josephine M. Pemberton

**Affiliations:** Institute of Evolutionary Biology, School of Biological Sciences, University of Edinburgh, Edinburgh, UK, EH9 3FL; Department of Biology, Indiana University, Bloomington, Indiana, 47405; Moredun Research Institute, Pentlands Science Park, Bush Loan, Midlothian, UK, EH26 0PZ

## Abstract

Spatial heterogeneity in parasite susceptibility and exposure is a common source of confounding variation in disease ecology studies. However, it is not known whether spatial autocorrelation acts on immunity in particular at small scales, within wild animal populations, and whether this predicts spatial patterns in infection. Here we used a well-mixed wild population of individually recognised red deer (*Cervus elaphus*) inhabiting a heterogeneous landscape to investigate fine-scale spatial patterns of immunity and parasitism. We noninvasively collected 842 faecal samples from 141 females with known ranging behaviour over two years. We quantified total and helminth-specific mucosal antibodies and counted propagules of three gastrointestinal helminth taxa. These data were analysed with linear mixed models using the Integrated Nested Laplace Approximation (INLA), using a Stochastic Partial Differentiation Equation approach (SPDE) to control for and quantify spatial autocorrelation. We also investigated whether spatial patterns of immunity and parasitism changed seasonally. We discovered substantial spatial heterogeneity in general and helminth-specific antibody levels and parasitism with two helminth taxa, all of which exhibited contrasting seasonal variation in their spatial patterns. Notably, strongyle nematode intensity did not align with density hotspots, while *Fasciola hepatica* intensity appeared to be strongly influenced by the presence of wet grazing. In addition, antibody hotspots did not correlate with distributions of any parasites. Our results suggest spatial heterogeneity may be an important factor affecting immunity and parasitism in a wide range of study systems. We discuss these findings with regards to the design of sampling regimes and public health interventions, and suggest that disease ecology studies investigate spatial heterogeneity more regularly to enhance their results, even when examining small geographic areas.

## Introduction

Parasite infection in the wild is extremely spatially heterogeneous. The scale at which spatial variation acts depends on the host and parasite being studied, and even fine-scale environmental heterogeneity may influence the spatial epidemiology of human diseases (Murdock *et al.* 2017). However, the spatial ecology of disease is most often considered in terms of large-scale patterns (e.g. Murray *et al.*, 2018) and using occurrence or prevalence data, which is less informative than intensity, particularly for macroparasites. In addition, due to practical considerations, many studies investigating spatial variation in wild animals compare several discrete populations rather than sampling across a continuous, mixing population (e.g. Downs *et al.* 2015; Cheynel *et al.* 2017). Alternatively, some studies rely on opportunistic convenience sampling, which can produce an inaccurate representation of disease processes and bias estimates of infection prevalence due to their non-random sampling in space (Nusser *et al.* 2008). As a result, little is known about fine-scale patterns of susceptibility and exposure, and how they influence spatial patterns of infection, in wild animals.

Identifying the relevant spatial scale for disease processes such as susceptibility and exposure is important, as quantifying spatial trends at different scales can introduce uncertainty at best, and can profoundly affect the conclusions drawn at worst (Gilligan *et al.* 2007; Vidal-Martínez *et al.* 2010; Lachish and Murray 2018). For example, Lyme disease risk correlates positively with biodiversity at the within-forest level, but the reverse is true between forests (Wood and Lafferty 2013). An understanding of spatial processes is therefore crucial for designing public health interventions (Caprarelli and Fletcher 2014) and sampling regimes (Nusser *et al.* 2008; Vidal-Martínez *et al.* 2010; Lachish and Murray 2018). A deeper understanding of fine-scale spatial variation in disease processes could also inform patterns seen over wider distances (Murdock *et al.* 2017; Pawley and McArdle 2018). In addition, if immunity and parasitism vary over short distances, infection-oriented studies of wild populations could be affected by greater degrees of spatial dependence than previously considered, which can affect inference. When spatial autocorrelation is not considered, the type I error rate may be inflated due to inflated covariance of explanatory and/or response variables emerging from geographic proximity (Pawley and McArdle 2018).

Spatial variation in immunity can originate from gradients in abiotic conditions such as temperature (Laughton *et al.* 2017) or in biotic factors such as prey availability (Becker *et al.* 2018). Spatial variation in parasitism will arise in part as a result of this immune heterogeneity owing to variation in susceptibility, clearance, and tolerance (Jolles *et al.* 2015), as well as from abiotic factors affecting parasite transmission (e.g. sunlight; Parsons *et al.* 2015) or from variation in abundance of secondary hosts or vectors (Sol *et al.* 2011; Olsen *et al.* 2015). In addition, conspecific density can influence resource availability, immune investment, and parasite exposure (Wilson *et al.* 2004; Downs *et al.* 2015; Ezenwa *et al.* 2016). We therefore expect to see considerable spatial variation in both immunity and parasitism in heterogeneous environments; where gradients are steep and mixing is minimal, this variation should occur over short distances. A recent study in wild mice (*Mus musculus*) demonstrated high between-site immune heterogeneity, but with extensive variation in the degree of within-site differentiation, suggesting short-range spatial dependence (Abolins *et al.* 2018). However, few studies have examined how both immunity and parasitism vary continuously across space within wild animal populations, so it is unclear to what degree spatial variation in parasitism in the wild originates from immune-mediated processes rather than from environmental factors affecting exposure. Finally, spatial patterns are rarely static, and may change over time (Hawkins 2012), yet seasonal or annual changes in these spatial patterns are rarely examined.

The red deer (*Cervus elaphus*) is a large land mammal closely related to the American wapiti (*Cervus canadensis*) whose distribution covers much of Europe. The relationship between red deer disease and their spatial behaviour is important to pathogen spillover, as this species carries a plethora of parasites that can infect humans and livestock (Bohm *et al.* 2007; Brites-Neto *et al.* 2015) and which they can vector between farms and distribute through the landscape (Chintoan-Uta *et al.* 2014; Qviller *et al.* 2016). The wild red deer living in the North Block of the Isle of Rum in Scotland are individually recognised and regularly censused, providing detailed information on each individual’s life history and ranging behaviour (Clutton-Brock *et al.* 1982). These censuses have previously been used to uncover important roles of the environment and spatial behaviour in influencing individuals’ phenotypes (Stopher *et al.* 2012; Froy *et al.* 2018). Longitudinal noninvasive faecal sampling of the population has revealed a high prevalence of several gastrointestinal helminth parasites including strongyle nematodes, the liver fluke *Fasciola hepatica*, and the tissue nematode *Elaphostrongylus cervi* (Albery, Kenyon, *et al.* 2018). The life cycle of strongyle nematodes is direct, while *F. hepatica* must infect *Galba truncatula* water snails (Taylor *et al.* 2016) and *E. cervi* infects a range of land snails (Mason 1989). Their mucosal antibodies (IgA) have also been quantified by faecal ELISA, offering a measure of immune investment (Albery, Watt, *et al.* 2018). Both helminth intensity and IgA concentrations are affected by deer reproductive investment and fluctuate seasonally (Albery, Watt, *et al.* 2018). However, the spatial distributions of these immune and parasite measures have yet to be investigated.

In this study, we used regular census data and noninvasive faecal samples from the deer population to investigate how individuals’ spatial behaviour was associated with immunity and parasitism at fine spatial scales. We incorporated spatial autocorrelation structures in order to investigate how this affected model fit, to identify hotspots of immunity and infection, and to quantify the spatial scale at which our data were autocorrelated. We also allowed spatial autocorrelation structures to vary seasonally. We expected that accounting for spatial autocorrelation would improve model fit, and that this would be a more effective way of investigating spatial trends than separating the population into discrete arbitrary subpopulations as was previously done to control for spatial variation (Huisman et al. 2016). We also predicted that individuals living in different areas of the study system would exhibit notably different antibody levels and parasite intensities. In particular, we expected that higher deer density would correspond to areas characterized by weaker immunity and in turn higher parasite intensity, as high density will correspond to higher competition for resources and increased parasite exposure (Wilson *et al.* 2004). Finally, we predicted that *F. hepatica* and *E. cervi* intensity would be influenced by the habitats of their secondary hosts – particularly that *F. hepatica* would be more common in wetter areas (Olsen *et al.* 2015).

## Methods

### Study system and sampling regime

The study population is located in the north block of the Isle of Rum, Scotland (57°N, 6°20′W; Figure 1). The sampling area measures ~4 km north-south and ~3 km west-east (total area ~12.7km^2^).The most intensely sampled area consists of a river running from south to north along a valley, flanked by hills on either side, and an extended ranging area around the coast to the east, close to the sea. Peat bogs and *Juncus* marshland comprise much of the southern and central areas of the valley, while the hills are dominated by wet and dry heath and *Molinia* grassland. In the north, moving seaward, the landscape is dominated by *Agrostis* and *Festuca* grassland, followed by sandy dunes and beaches. The study population is wild and unmanaged, and is censused five times a month for eight months of the year (see Clutton-Brock *et al.*, 1982). During censusing, one of two predetermined routes is walked or driven through the study area and individuals’ locations (to the nearest 100 metres) are noted. The northern part of the study area hosts the highest population density, with most deer centred around the high-quality grazing near the mouth of the river and the land around the coast to the east (Figure 1). Annual home ranges are highly repeatable from year to year (Stopher *et al.* 2012).

**Figure 1:**
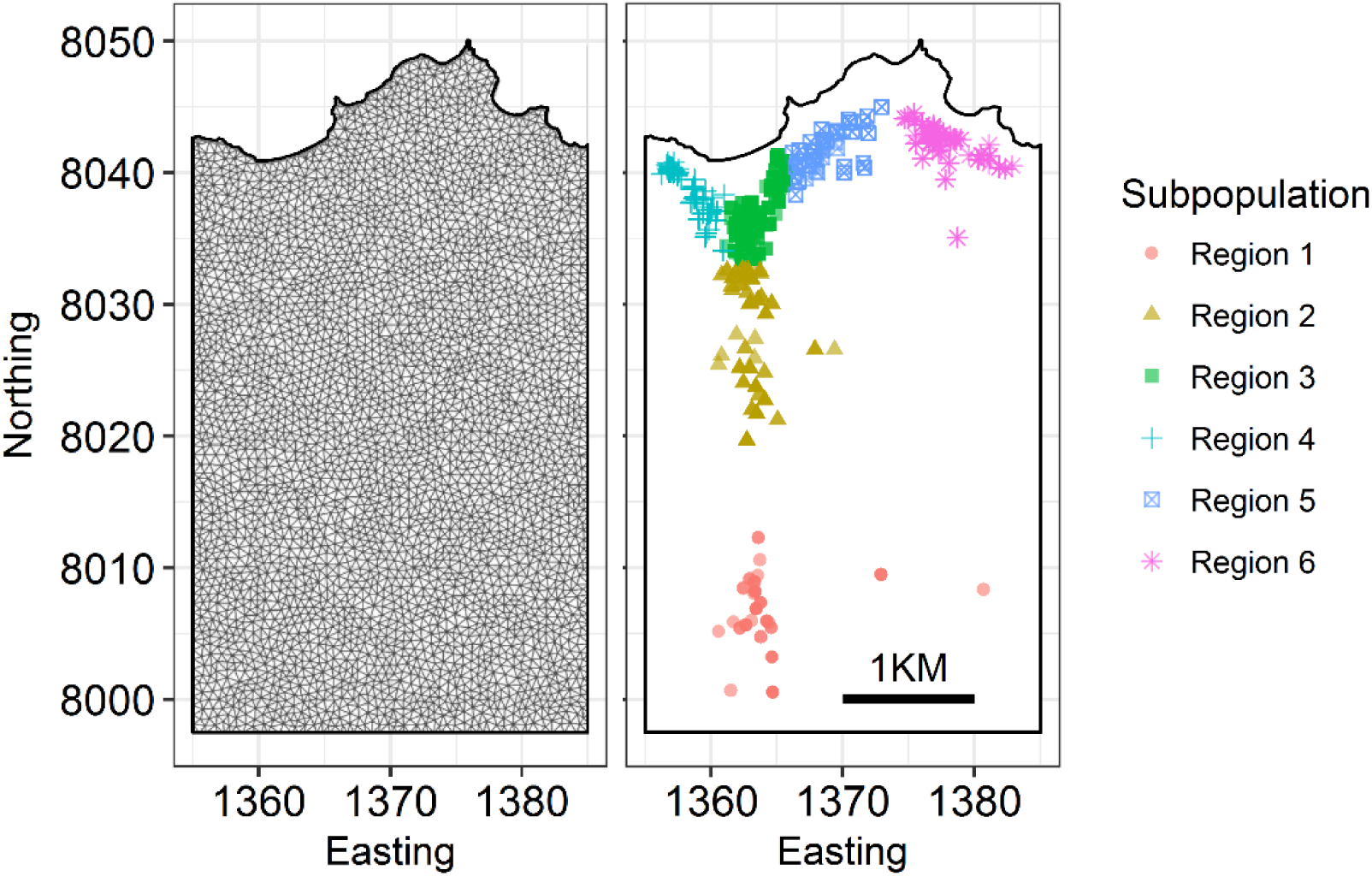
Map of the Isle of Rum red deer study area, depicting the mesh used for the INLA SPDE random effect (left) and the sampling locations and their subpopulations (right). Easting and Northing are in units of 100m grid squares, so that 10 units corresponds to 1km, per the scale bar. the river at the base of the valley runs south-north along the 1363 Easting. Subpopulations are organised and named running from south to north, then west to east.

The deer reproductive cycle (“deer year”) spans from the start of the calving season, May 1^st^, until April 30^th^ the following year. Samples were collected as previously described (Albery, Kenyon, *et al.* 2018), on a seasonal basis during 7 two-week trips in August (“summer”), November (“autumn”) and April (“spring”) between April 2016 and April 2018 inclusive. Note that our dataset included a sampling trip from April 2016, which was part of the deer year beginning in May 2015, with no accompanying summer and autumn trips from this reproductive cycle. In the study period, 842 faecal samples were collected noninvasively from 141 individually known adult females aged 3 and above. Parasite propagule counts and antibody ELISA quantification were carried out on these samples as previously described (Albery, Kenyon, *et al.* 2018; Albery, Watt, *et al.* 2018). Parasites included strongyle nematodes (order: Strongylida), the common liver fluke *Fasciola hepatica* and the red deer tissue nematode *Elaphostrongylus cervi*. Our two antibody measures were total mucosal IgA levels (“total IgA”) and anti-*Teladorsagia circumcincta* L3 larval antigen IgA (“anti-Tc IgA”). The former is taken as an indicator of general investment in mucosal immunity, while the latter gives a measure of specific anti-strongyle IgA response which is thought to be more indicative of protective immunity against strongyles (Watt *et al.* 2016; Albery, Watt, *et al.* 2018). There was not enough faecal matter in all samples to quantify all variables; final sample sizes are displayed in Table 1. Using the census data, each individual’s mean easting and northing over the deer year was taken as their average location. This was taken to be a better indication of an individual’s spatial behaviour than the location at which the faecal sample itself was collected. We subdivided the study area into six approximate subpopulations based on each individual’s average location (Huisman *et al.* 2016). These locations and subpopulations are displayed in Figure 1.

**Table 1:** Model set descriptions and extracted values (ΔDIC, halving range, ρ and sample size) for each response variable (Str = strongyles; Fh = *F. hepatica*; Ec = *E. cervi*; TotA = Total IgA; TcA = anti-Tc IgA). The model with the lowest DIC for each response variable (ΔDIC=0) is highlighted in bold and underlined; where the best-fitting models are not distinguishable (ΔDIC<2), both are highlighted and underlined. The halving range of each variable represents the distance in metres at which spatial autocorrelation reduces to 0.5, either for model set 3 or for the best-fitting seasonally varying model (from model set 5, or model set 4 for total IgA). ρ value estimates are given with their 0.025 and 0.975 quantiles in brackets.

| Model Set | Description | Str | Fh | Ec | TotA | TcA |
| --- | --- | --- | --- | --- | --- | --- |
| **ΔDIC** |  |  |  |  |  |  |
| 1 | Base set | 130.13 | 8.27 | <u>0</u> | 39.27 | 22.81 |
| 2 | + Subpopulation fixed effect | 132.13 | 6.49 | 3.78 | 37.8 | 26.78 |
| 3 | + Spatial field random effect | 102.42 | 4.25 | 1.36 | 32.33 | 19.37 |
| 4 | + Field varying seasonally | <u>0</u> | 2.29 | 6.11 | <u>0</u> | 2.02 |
| 5 | + correlation between fields | 4.08 | <u>0</u> | 2.26 | <u>1.08</u> | <u>0</u> |
| **Autocorrelation halving ranges (metres)** |  |  |  |  |  |  |
| 3 | Static spatial field | 59.62 | 1323.58 |  | 150.06 | 92.58 |
| 4-5 | Seasonally varying spatial field | 32.58 | 1124.74 |  | 640.07 | 415.82 |
| **$\rho$ (correlation between seasonal fields)** |  |  |  |  |  |  |
| 5 | Seasonally varying spatial field | 0.09<br>(-0.24,0.45) | 0.67<br>(-0.02,0.98) |  | -0.1<br>(-0.44,0.48) | -0.31<br>(-0.5,0.3) |
| **Sample Size** |  |  |  |  |  |  |
| 1-5 | Number of samples | 834 | 823 | 832 | 799 | 795 |
| 1-5 | Number of individuals | 139 | 139 | 138 | 137 | 137 |

### Statistical analysis

Statistical analysis was carried out using the Integrated Nested Laplace Approximation (INLA). INLA is a deterministic Bayesian approach which is increasingly being used for analysis of spatial data (Zuur *et al.* 2017). Models were fitted in R version 3.5 (R Core Team 2018) using the linear modelling package R-INLA (Rue and Martino 2009; Martins *et al.* 2013). We constructed 5 generalised linear mixed models (GLMMs) for each response variable, each featuring different combinations of fixed and spatial random effects. The distinguishing components of these model sets are outlined below and displayed in Table 1.

Our five response variables included integer counts per gram of three parasite propagules following a negative binomial distribution (strongyles, *F. hepatica* and *E. cervi*) and Gaussian-distributed optical densities of two mucosal antibodies (total IgA and anti-Tc IgA). Antibody levels were corrected for collection effects as previously described, by taking the residuals from a linear model including raw antibody OD as a response variable and including day of collection, time of collection and extraction session as explanatory variables (Albery, Watt, *et al.* 2018). In our main GLMMs, explanatory variables included: Deer year (categorical with three levels: 2015, 2016, and 2017); Season (categorical with three levels: Summer, Autumn, and Spring); Age (continuous, in years); Reproductive status (categorical variable with three levels: No Calf, Calf Died, and Calf Survived; see Albery, Watt, *et al.* (2018) for definitions); Subpopulation (categorical, six levels). All models included individual ID as a random effect.

INLA allows incorporation of a spatially distributed random effect to account for spatial autocorrelation (Lindgren *et al.* 2011). This uses a stochastic partial differentiation equation (SPDE) approach to approximate the continuous random field using a triangulated mesh of connected discrete locations (Lindgren and Rue 2015). The mesh we used for the spatial random effect is displayed in Figure 1. The random effect can be plotted in 2D (giving the “spatial field”) to investigate hot-and coldspots of the response variable, and the kappa/range parameters can be extracted to investigate the distance at which autocorrelation fades in space. It is also possible to allow multiple spatial fields within a single model, by assigning separate fields to different categories or by linking fields with correlation structures to investigate spatiotemporal variation. The underlying mathematics of INLA and associated spatial/spatiotemporal models have been extensively discussed elsewhere, and such models are increasingly being used to examine spatiotemporal trends (e.g. fisheries ecology (Cosandey-Godin *et al.* 2015)); see http://www.r-inla.org for more examples.

We constructed a set of competing models for each response variable. Each model set contained five models, resulting in 25 models total. Our base model set (model set 1) included year, season, age, and reproductive status as fixed effects, similar to models previously used to investigate associations between reproduction, immunity, and parasitism (Albery, Watt, *et al.* 2018). Model set 2 added subpopulation as a fixed effect to investigate whether this explained any variation and to examine the value of analysing continuous populations using discrete subdivisions (Figure 1). Model set 3 added a spatially distributed SPDE random effect, rather than the subpopulation fixed effect, to control for and quantify spatial autocorrelation. In model set 4, this spatial field was allowed to vary between seasons (summer, autumn, and spring), and model set 5 allowed correlation between these seasonal fields. To allow spatial fields to correlate, we used an “exchangeable” model, where all fields in the model were correlated by the same value (ρ) rather than e.g. following an autoregressive process through time.

For each response variable, the five fitted models were compared using Deviance Information Criterion (DIC). A change in 2 DIC was selected to distinguish between models and select the most parsimonious model. When the best-fitting models included spatial autocorrelation, we extracted the range parameters to estimate the range of autocorrelation and ρ parameters to estimate correlation between seasonal fields. For the range of autocorrelation, we report the distance at which spatial autocorrelation decayed to 0.5 (henceforth “halving range”; (Brooker *et al.* 2006)). Finally, we compared effect sizes from each model to investigate whether incorporating spatial autocorrelation altered any conclusions about the fixed effects. We particularly focussed on whether accounting for spatial autocorrelation altered the estimates for reproductive status effects, which have previously been demonstrated to impact both immunity and parasitism, and vary spatially across the population.

## Results

Our models revealed strong and contrasting spatial trends in all but one of our response variables. All models but *E. cervi* were incrementally improved by first incorporating a spatial random effect and then by allowing it to vary between seasons (DIC values in Table 1; all secondary models had ΔDIC≥3.44). In all cases, including spatially distributed random effects improved model fit compared to fitting a subpopulation fixed effect (Table 1; ΔDIC≥2.4). The spatial fields of the random effects, taken from model sets 3-5, are displayed in Figure 2. For each response variable, we report the spatial field and results from both model set 3 (spatial field constant across the study period) and model set 4 (spatial seasons varying seasonally, with no correlation between fields). The exception is *F. hepatica*, for which allowing the seasonal fields to correlate in model set 5 improved model fit (ΔDIC=-2.29, Table 1); therefore, for *F. hepatica,* we display the fields and results from model sets 3 and 5. Response variables differed considerably in terms of both their spatial fields (Figure 2) and the range at which they varied (Figure 3). Table 1 also displays halving ranges and ρ values; as *E. cervi* models were never improved by the inclusion of the subpopulation fixed effect or by SPDE random effects (ΔDIC>1.36), we do not report these results further.

**Figure 2:**
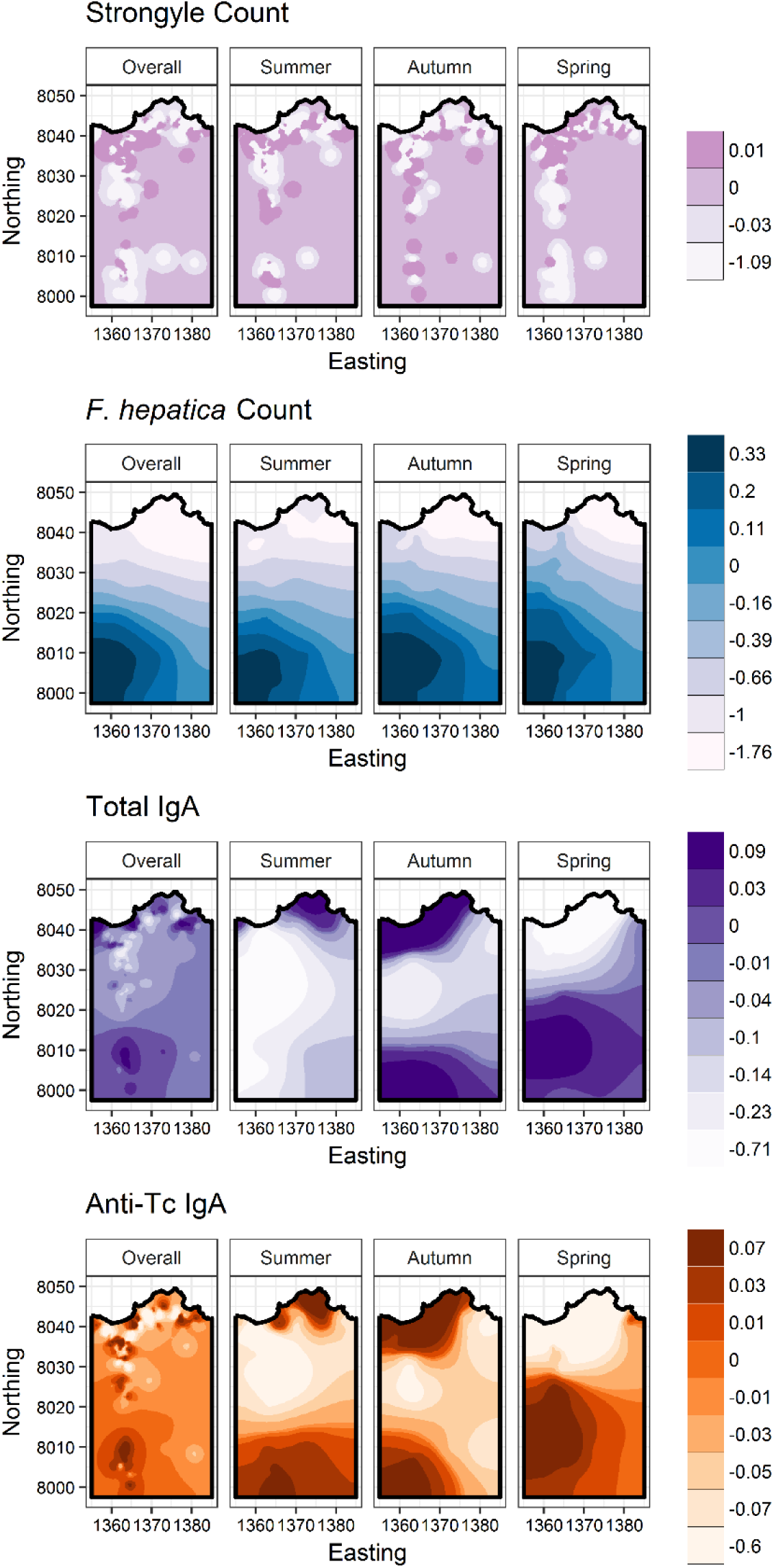
Projections of the spatially distributed SPDE random effect (spatial fields) for the models that were improved by its inclusion (one row for each variable, from top row to bottom: strongyles; *F. hepatica*; total IgA; anti-Tc IgA). Spatial fields were taken from model sets 3 (constant spatial field, far left column) and 4-5 (spatial fields varying seasonally, remaining three columns). Colours denote the lower bounds of 9 quantiles of the spatial effects on the link scale, rounded to 2 decimal places, with darker colours representing higher parasite counts (rows 1-2) or antibody levels (rows 3-4). Where there are fewer than 9 colours, this is because rounding the values to 2 decimal places created identical quantile values, and demonstrates that spatial autocorrelation accounted for a smaller proportion of the variation. Easting and Northing are in units of 100m grid squares, so that 10 units corresponds to 1km. the river at the base of the valley runs along the 1363 Easting.

**Figure 3:**
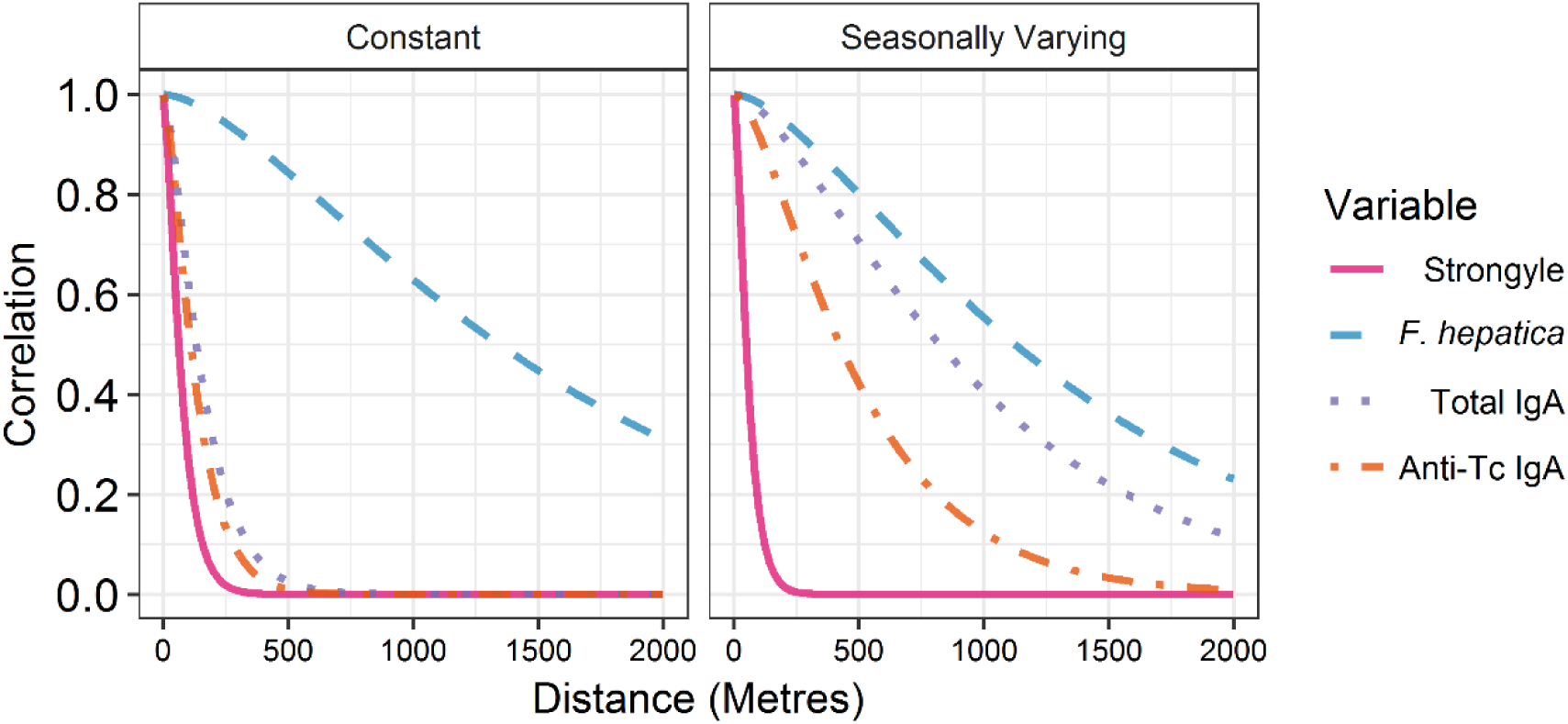
the range of spatial autocorrelation acting on each response variable, in metres, for INLA models with constant spatial fields (model set 3, left) and those with spatial fields that varied between seasons (model sets 4-5, right). In the right panel, values were taken from the best-fitting seasonally varying model: model 4 for total IgA, and model 5 for the remaining response variables (Table 1). See Table 1 for the halving ranges for each variable when the field was kept constant or allowed to vary seasonally.

Strongyle nematode intensity exhibited weak spatial patterns, with a very short range of autocorrelation; this did not increase when spatial fields were allowed to vary seasonally (Figure 2-3, halving range<59.62M). Allowing the spatial field to vary between seasons resulted in similar patchy distributions which are hard to distinguish (Figure 2) but nevertheless improved model fit compared to all other models (Table 1, ΔDIC=4.25). *F. hepatica* demonstrated a strong spatial pattern, with high intensities in the mid-and south-valley decreasing to the north and northeast (Figure 2). This gradual, unidirectional trend was reflected in the long range of autocorrelation (Figure 3, halving range=1323M). Allowing the spatial field to vary between seasons improved *F. hepatica* model fit, but resulted in similar seasonal fields (Figure 2). This was reflected in the positive ρ parameter (ρ=0.67) derived from model 5, which was the best-fitting model for *F. hepatica*, demonstrating that seasonal spatial fields were substantially positively correlated.

When the spatial field was kept constant across the study period, total IgA and anti-Tc IgA both demonstrated a very short range of spatial autocorrelation (Figure 3, halving range<150.06M). Both antibody distributions were similar and negatively correlated with that of strongyles, being lower in the central north and higher in the south and edges of the study area (Figure 2). However, allowing both antibodies’ spatial fields to vary between seasons improved model fit substantially compared to all other models (Table 1, ΔDIC<18.58), increased the range of autocorrelation (Figure 3, halving range>415.82M), and resulted in very different seasonal patterns (Figure 2). These patterns were similar between total IgA and anti-Tc IgA, although total IgA had a slightly larger range of autocorrelation (Figure 3, halving range=640.07M and 415.82M for total IgA and anti-Tc IgA respectively). The best-fitting model for total IgA and anti-Tc IgA was either model 4 or 5 for total IgA (Table 1, ΔDIC<2), while model 5 fit slightly better for anti-Tc IgA (ΔDIC=2.02). Hence model 4 is presented for total IgA as the model with fewer degrees of freedom, and model 5 is presented for anti-Tc IgA.

The subpopulation fixed effects broadly followed the spatial fields of the SPDE random effects (Figure 4). Briefly, strongyles showed little difference across different regions, although estimates for the two northern regions (regions 3 and 5) did not overlap with zero when compared to the southern region 1. For *F. hepatica* intensities decreased moving northeast from region 1 to region 6, and all regions exhibited significantly decreased levels below the far south region 1. The reverse was true for *E. cervi* intensities. Patterns for total IgA and anti-Tc IgA are harder to interpret and less significant, but broadly the far south region 1 subpopulation featured higher antibody levels than northern regions (regions 2, 3, and 5 for total IgA and region 5 for anti-Tc IgA).

**Figure 4:**
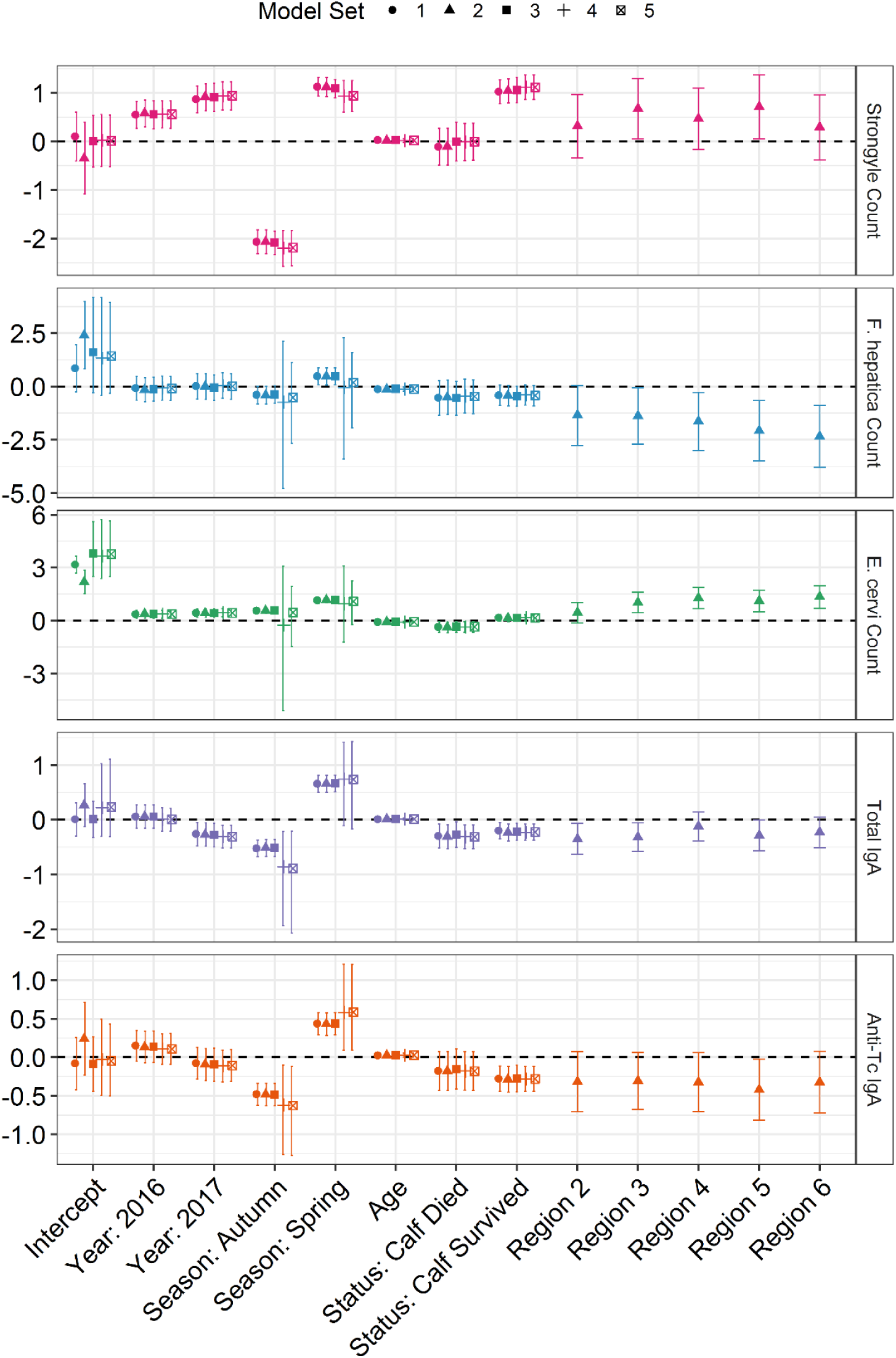
Comparison of the fixed effect estimates from each model for each response variable. Points denote the mean effect estimate, and error bars represent the 0.025 and 0.975 quantiles. Estimates for categorical variables represent departures from the missing category of each variable (year: 2015, season: summer, status: no calf, subpopulation: region 1), on the link scale. Points of different shapes denote the results from different model sets. Model set 1: base model. Model set 2: base model + subpopulation fixed effect. Model set 3: base model + spatial random effect. Model set 4: spatial random effect allowed to vary between seasons. Model set 5: spatial random effect allowed to correlate between seasons.

Most fixed effect estimates were only slightly modified by incorporating spatial autocorrelation structures in our models (Figure 4). No estimates were reduced in significance except the seasonal effects in models 4 and 5 for *F. hepatica* and *E. cervi* (Figure 4). Examining the spatial fields (Figure 2), this reduction in seasonal effect probably originated from competition between the seasonally varying spatial random effect and the season variable itself. Otherwise, effect estimates remained unchanged when spatial autocorrelation was included. This was particularly true for reproductive status effects, many of which actually increased slightly in magnitude when we accounted for spatial autocorrelation (Figure 4).

## Discussion

This study has revealed fine-scale spatial variation in immunity and parasitism at an individual level in a large wild mammal population. Spatial heterogeneity contributed considerably to between-individual differences in immunity and parasitism despite a total sampling area of only ~12.7 km^2^. The scale of spatial dependence was therefore extremely short, and well within the scale of the study area. These findings are in accordance with a previous study demonstrating fine-scale immune variation in a discrete spatial context (within-site versus between-site) in wild mice (Abolins *et al.* 2018). We demonstrate similar spatial variation in a continuous context, and in both antibody levels and parasite counts, despite considerable mixing within the population. Furthermore, the response variables differed in terms of their spatial fields, the range at which spatial dependence acted, and their interactions with seasonality. Finally, spatial distributions of antibodies were not similar to any parasite distributions, implying that fine-scale environmental factors acting on exposure are more important than host immune susceptibility in driving spatial heterogeneity of parasite infection.

### The scale of dependence and its importance for disease ecology studies

Understanding the spatiotemporal scale of disease processes is important for designing sampling regimes and disease control strategies (Caprarelli and Fletcher 2014; Lachish and Murray 2018). In this context, our results have several important general implications. Firstly, fine-scale trends like those exhibited here may scale up quickly, contributing to larger-scale geographic patterns of disease that are more commonly studied (Ostfeld *et al.* 2005; Murdock *et al.* 2017; Murray *et al.* 2018). Second, disease ecology studies that do not consider spatial autocorrelation, even over short distances, may be missing important sources of variation in immunity and exposure and risk reporting biased effect estimates. The persistent spatial trend seen in *F. hepatica* (Figure 2) demonstrates that different areas of a given study system can be consistently associated with either higher or lower parasitism, so that uneven sampling in space could introduce confounding variation and bias. In contrast, where the range of autocorrelation is extremely short, as in strongyles, sampling regimes that do not consider spatial dependence may incidentally sample areas of both high and low parasitism, reducing the risk of spatial biasing. Trends are not necessarily similar across variables, complicating matters: most notably, spatial gradients of *F. hepatica* and strongyle intensity differed considerably both in range and patterns, and antibody hotspots did not align with parasite hotspots (Figure 2). Therefore, information on the spatial distribution of one immune or parasite measure could not be used to infer the distribution of another, and appropriate sampling regimes will differ between response variables. Finally, all models except *E. cervi* were further improved when the spatial field was allowed to vary temporally, and spatial patterns of antibody levels changed considerably between seasons (Figure 2). This confirmed our expectations that spatial fields would not be static in time (Hawkins 2012). Therefore, in some cases, even sampling from a wide, contiguous area may only capture a cross-sectional snapshot of the spatial dynamics of a given study system, necessitating longitudinal analysis.

Spatial heterogeneity has the potential to obscure or produce artefactual associations with other variables, modifying conclusions drawn from models without spatial dependence structures – in particular by inflating the type I error rate (Beale *et al.*, 2010; Pawley and McArdle, 2018). However, in this study, fixed effects remained largely unchanged when incorporating spatial dependence structures despite the importance of spatial heterogeneity (Figure 4). In particular, previously reported reproductive status effects (Albery, Watt, *et al.* 2018) persisted or increased slightly in size, despite the fact that reproductive success varies across the study area (McLoughlin *et al.* 2006; Stopher *et al.* 2012). This demonstrates that spatial variation can contribute to ecological patterns of disease without necessarily obscuring other findings (Pawley and McArdle 2018). We suggest that disease ecology studies that involve wild sampling attempt to investigate spatial variation to enrich their results, rather than viewing spatial autocorrelation as a nuisance (Pawley and McArdle 2018). In addition, although the spatial fields were broadly reflected by the subpopulation fixed effect results (Figure 4), the spatial fields were more easily interpretable and increased model fit, and therefore incorporating spatial autocorrelation was advantageous. While integrating spatial dependence did not have severe impacts on effect sizes in our study, we lastly encourage researchers to consider accounting for spatial dependence even at the fine scales here to improve statistical inference and account for this variation.

### Interpreting the spatial fields

The spatial fields derived from our models can help to indicate the factors determining immunity and parasite infection. Spatial trends of *F. hepatica* were especially stark, being much higher in the south of the study area and decreasing to the north and northeast (Figure 2). Given that the parasite distributions were not explained through differences in immune susceptibility, particularly considering minimal overlap with antibody level distributions (Figure 2), spatial patterns in parasite intensity likely instead resulted from spatial variation in exposure. This heterogeneity likely originated from the drier environment in the north compared to the wet, marshy ground in the south of the valley, the latter of which could be conducive to parasite persistence in the environment. After being excreted, *F. hepatica* eggs develop to form infectious miracidia, which seek out and infect *Galba truncatula* water snails (Taylor *et al.* 2016; Beesley *et al.* 2018). After a period within the snail, cercariae are produced which encyst on vegetation as metacercariae to be consumed by deer. Wet areas are likely to host higher *G. truncatula* abundance, and warmer, wetter environments are conducive to fluke development and host seeking behaviour, both of which will produce higher exposure (Ollerenshaw and Smith 1969). The observed fluke distribution agrees with a number of studies in livestock demonstrating high fluke risk where grazing and wet areas intersect (e.g. Olsen *et al.*, 2015). Similar relationships with water sources are displayed by the human trematode *Schistosoma mansoni*, which shows a similar range of autocorrelation (Brooker *et al.* 2006). Our corroboration of these findings in a wild mammal implies that similar environmental risk factors may be influencing trematode infection in wild animals, humans, and livestock.

In contrast to *F. hepatica*, the spatial field of strongyle intensity is difficult to interpret: spatial autocorrelation introduced important variation, yet the range of autocorrelation was small, with no discernible pattern either across the study period nor within seasons (Figure 2). Strongyles’ very small autocorrelation range was similar to that reported for human hookworm infection (Brooker *et al.* 2006), but surprisingly the overall pattern goes against our expectation that strongyle exposure (and therefore infection intensity) should correlate with local deer density (Wilson *et al.* 2004). Strongyles may be less impacted by environmental factors than are *F. hepatica* and *E. cervi* due to their direct life cycle, which does not involve a secondary host, such that spatial autocorrelation in intrinsic factors affecting susceptibility is more important than environmental effects on exposure and transmission. Host genetic similarity is one possible intrinsic factor producing the spatial autocorrelation seen in strongyle counts and antibody levels: both are heritable in ungulates (Bisset *et al.* 1992; Callaby *et al.* 2014; Hayward *et al.* 2014), and genetic relatedness is correlated with spatial distance in this system (Stopher *et al.* 2012).

### Ecological and epidemiological implications

The fine-scale spatial heterogeneity demonstrated here has implications for the ecology and control of infectious disease in wild ungulate populations. For example, localised transmission hotspots may maintain parasite diversity, preventing parasite strains and species from outcompeting each other through geographic niche differentiation and contributing to the considerable genetic differentiation seen in liver fluke populations (Beesley *et al.* 2016). Additionally, when combined with sex-specific deer ranging patterns, spatial trends could contribute to previously observed sex biases (Albery, Kenyon, *et al.* 2018).

As *F. hepatica* is an important livestock parasite, fluke control initiatives should consider the presence of high-risk wet areas of grazing that may be used by deer populations. However, it is worth noting that the fluke hotspots here were observed at the per-capita count level, rather than as an absolute number of parasites in the environment. Given the higher deer density in the north, taking *F. hepatica* as an example, it is likely that the absolute number of fluke eggs being excreted in the north is higher than the south, but these parasites are less likely to complete their life cycle due to unsuitable environmental conditions. In the future, it may be possible to compare the excretion and movement patterns of the deer with pasture larval counts and snail sampling across the study area to examine the rate at which successful infection occurs, and to investigate whether deer living in the high-risk southern area of the valley may indeed be vectoring *F. hepatica* to the north (Chintoan-Uta *et al.* 2014; French *et al.* 2016).

## Author contributions

GFA designed the study, collected samples, conducted labwork, analysed the data, and wrote the manuscript. DJB, FK, DHN, and JMP offered comments and suggestions on theory, analysis, and the manuscript throughout.

## Acknowledgements

The long term red deer study is funded by the Natural Environment Research Council (grant number NE/L00688X/1), as is GFA’s PhD studentship through the E3 Doctoral Training Partnership (grant number NE/L002558/1). FK receives funding from the Scottish Government, RESAS, Strategic Research Programmes 2016-21. We thank Scottish Natural Heritage for permission to work on the Isle of Rum NNR and for the support of the reserve management team on the island. Thanks to Dave McBean and Gillian Mitchell at the Moredun Research Institute for their help with parasitological methods. The *Teladorsagia circumcincta* antigen was received from Moredun Research Institute, and was prepared by David Bartley, Alison Morrison, Leigh Andrews, David Frew and Tom McNeilly. Thanks also to Kathryn Watt for assisting with immunological labwork and to Sean Morris, Alison Morris, Olly Gibb, and all other field assistants for their help in sample collection.

## References

Abolins S, Lazarou L, Weldon L, Hughes L, King EC, Drescher P, Pocock MJO, Hafalla JCR, Riley EM, Viney M. 2018. The ecology of immune state in a wild mammal, Mus musculus domesticus. PLOS Biol 16:e2003538.

Albery GF, Kenyon F, Morris A, Morris S, Nussey DH, Pemberton JM. 2018. Seasonality of helminth infection in wild red deer varies between individuals and between parasite taxa. Parasitology 145:1–11.

Albery GF, Watt K, Keith R, Morris S, Morris A, Kenyon F, Nussey DH, Pemberton JM. 2018. Reproduction has different costs for immunity and parasitism in a wild mammal. bioRxiv 472597.

Beale CM, Lennon JJ, Yearsley JM, Brewer MJ, Elston DA. 2010. Regression analysis of spatial data. Ecol Lett 13:246–64.

Becker DJ, Czirják GÁ, Volokhov D V., Bentz AB, Carrera JE, Camus MS, Navara KJ, Chizhikov VE, Fenton MB, Simmons NB, Recuenco SE, Gilbert AT, Altizer S, Streicker DG. 2018. Livestock abundance predicts vampire bat demography, immune profiles, and bacterial infection risk. Philos Trans R Soc B in press: doi:10.1098/rstb.2017.0089.

Beesley NJ, Caminade C, Charlier J, Flynn RJ, Hodgkinson JE, Martinez-Moreno A, Martinez-Valladares M, Perez J, Rinaldi L, Williams DJL. 2018. Fasciola and fasciolosis in ruminants in Europe: Identifying research needs. Transbound Emerg Dis 65:199–216.

Beesley NJ, Williams DJLL, Paterson S, Hodgkinson J. 2016. Fasciola hepatica demonstrates high levels of genetic diversity, a lack of population structure and high gene flow: possible implications for drug resistance. Int J Parasitol 47:11–20.

Bisset SA, Vlassoff A, Morris CA, Southey BR, Baker RL, Parker AGH. 1992. Heritability of and genetic correlations among faecal egg counts and productivity traits in Romney sheep. New Zeal J Agric Res 35:51–58.

Bohm M, White PCL, Chambers J, Smith L, Hutchings MR. 2007. Wild deer as a source of infection for livestock and humans in the UK. Vet J 174:260–76.

Brites-Neto J, Maria Duarte Roncato K, Martins TF. 2015. Tick-borne infections in human and animal population worldwide. Vet World 8:301–15.

Brooker S, Alexander N, Geiger S, Moyeed RA, Stander J, Fleming F, Hotez PJ, Correa-Oliveira R, Bethony J. 2006. Contrasting patterns in the small-scale heterogeneity of human helminth infections in urban and rural environments in Brazil. Int J Parasitol 36:1143–51.

Callaby R, Hanotte O, Conradie van Wyk I, Kiara H, Toye P, Mbole-Kariuki MN, Jennings A, Thumbi SM, Coetzer JAW, de C Bronsvoort BM, Knott SA, Woolhouse MEJ, Kruuk LEB. 2014. Variation and covariation in strongyle infection in East African shorthorn zebu calves. Parasitology 142:1–13.

Caprarelli G, Fletcher S. 2014. A brief review of spatial analysis concepts and tools used for mapping, containment and risk modelling of infectious diseases and other illnesses. Parasitology 141:581–601.

Cheynel L, Lemaître JF, Gaillard JM, Rey B, Bourgoin G, Ferté H, Jégo M, Débias F, Pellerin M, Jacob L, Gilot-Fromont E. 2017. Immunosenescence patterns differ between populations but not between sexes in a long-lived mammal. Sci Rep 7:1–11.

Chintoan-Uta C, Morgan ER, Skuce PJ, Coles GC. 2014. Wild deer as potential vectors of anthelmintic-resistant abomasal nematodes between cattle and sheep farms. Proc R Soc B 281:20132985.

Clutton-Brock TH, Guinness FE, Albon SD. 1982. Red Deer: Behavior and Ecology of Two Sexes. University of Chicago Press.

Cosandey-Godin A, Krainski ET, Worm B, Flemming JM. 2015. Applying Bayesian spatiotemporal models to fisheries bycatch in the Canadian Arctic. Can J Fish Aquat Sci 72:186–97.

Downs CJ, Stewart KM, Dick BL. 2015. Investment in constitutive immune function by north American elk experimentally maintained at two different population densities. PLoS One 10:1–17.

Ezenwa VO, Ghai RR, McKay AF, Williams AE. 2016. Group living and pathogen infection revisited. Curr Opin Behav Sci 12:66–72.

French AS, Zadoks RN, Skuce PJ, Mitchell G, Gordon-Gibbs DK, Craine A, Shaw D, Gibb SW, Taggart MA. 2016. Prevalence of liver fluke (Fasciola hepatica) in wild red deer (Cervus elaphus): Coproantigen elisa is a practicable alternative to faecal egg counting for surveillance in remote populations. PLoS One 11:1–18.

Froy H, Börger L, Regan CE, Morris A, Morris S, Pilkington JG, Crawley MJ, Clutton-Brock TH, Pemberton JM, Nussey DH. 2018. Declining home range area predicts reduced late-life survival in two wild ungulate populations. Ecol Lett 21:1001–9.

Gilligan CA, Truscott JE, Stacey AJ. 2007. Impact of scale on the effectiveness of disease control strategies for epidemics with cryptic infection in a dynamical landscape: An example for a crop disease. J R Soc Interface 4:925–34.

Hawkins BA. 2012. Eight (and a half) deadly sins of spatial analysis. J Biogeogr 39:1–9.

Hayward AD, Garnier R, Watt KA, Pilkington JG, Bryan T, Matthews JB, Pemberton JM, Nussey DH, Andrea L, Hayward AD, Garnier R, Watt KA, Pilkington JG, Grenfell BT, Matthews JB, Pemberton JM, Nussey DH, Graham AL. 2014. Heritable, heterogeneous, and costly resistance of sheep against nematodes and potential feedbacks to epidemiological dynamics. Am Nat 184 Suppl:S58–76.

Huisman J, Kruuk LEB, Ellis PA, Clutton-Brock T, Pemberton JM. 2016. Inbreeding depression across the lifespan in a wild mammal population. Proc Natl Acad Sci 201518046.

Lachish S, Murray KA. 2018. The Certainty of Uncertainty: Potential Sources of Bias and Imprecision in Disease Ecology Studies. Front Vet Sci 5:1–14.

Laughton AM, O’Connor CO, Knell RJ. 2017. Responses to a warming world: Integrating life history, immune investment, and pathogen resistance in a model insect species. Ecol Evol 7:9699–9710.

Lindgren F, Rue H. 2015. Bayesian Spatial Modelling with R-INLA. J Stat Softw 63. Lindgren F, Rue H, Lindstrom J. 2011. An explicit link between Gaussian fields and Gaussian Markov random fields?: the stochastic. 423–98.

Martins TG, Simpson D, Lindgren F, Rue H. 2013. Bayesian computing with INLA: New features. Comput Stat Data Anal 67:68–83.

Mason P. 1989. Elaphostrongylus cervi - a review. Surveillance 16:3–10.

McLoughlin PD, Boyce MS, Coulson T, Clutton-Brock T. 2006. Lifetime reproductive success and density-dependent, multi-variable resource selection. Proc R Soc B Biol Sci 273:1449–54.

Murdock C, Evans M V, McClanahan T, Miazgowicz K, Tesla B. 2017. Fine-scale variation in microclimate across an urban landscape changes the capacity of Aedes albopictus to vector arbovirus. PLoS Negl Trop Dis 11:e0005640.

Murray K, Olivero J, Roche B, Tiedt S, Guégan J. 2018. Pathogeography: leveraging the biogeography of human infectious diseases for global health management. Ecography (Cop) 1–17.

Nusser SM, Clark WR, Otis DL, Huang L. 2008. Sampling Considerations for Disease Surveillance in Wildlife Populations. J Wildl Manage 72:52–60.

Ollerenshaw CB, Smith LP. 1969. Meteorological Factors and Forecasts of Helminthic Disease. Adv Parasitol 7:283–323.

Olsen A, Frankena K, Bødker R, Toft N, Thamsborg SM, Enemark HL, Halasa T. 2015. Prevalence, risk factors and spatial analysis of liver fluke infections in Danish cattle herds. Parasit Vectors 8:160.

Ostfeld RS, Glass GE, Keesing F. 2005. Spatial epidemiology: An emerging (or re-emerging) discipline. Trends Ecol Evol 20:328–36.

Parsons SK, Bull CM, Gordon DM. 2015. Spatial variation and survival of Salmonella enterica subspecies in a population of australian sleepy lizards (Tiliqua rugosa). Appl Environ Microbiol 81:5804–11.

Pawley MDM, McArdle BH. 2018. Spatial autocorrelation: Bane or Bonus? bioRxiv 385526.

Qviller L, Viljugrein H, Loe LE, Meisingset EL, Mysterud A. 2016. The influence of red deer space use on the distribution of Ixodes ricinus ticks in the landscape. Parasit Vectors 9:545.

R Core Team. 2018. R: A language and environment for statistical computing. R Foundation for Statistical Computing, Vienna, Austria‥

Rue H, Martino S. 2009. Approximate Bayesian inference for latent Gaussian models by using integrated nested Laplace approximations. 319–92.

Sol D, Jovani R, Torres J, Sol D, Jovani R, Torres J. 2011. Geographical Variation in Blood Parasites in Feral Pigeons?: The Role of Vectors Geographical variation in blood parasites in feral pigeons?: the role of vectors. 23:307–14.

Stopher K V, Walling CA, Morris A, Guinness FE, Clutton-brock TH, Pemberton JM, Nussey DH. 2012. Shared spatial effects on quantitative genetic parameters: accounting for spatial autocorrelation and home range overlap reduces estimates of heritability in wild red deer. Evolution (N Y) 66:2411–26.

Taylor MA, Coop RL, Wall RL. 2016. Parasites of ungulates. In: Veterinary Parasitology p. 761–815.

Vidal-Martínez VM, Pech D, Sures B, Purucker ST, Poulin R. 2010. Can parasites really reveal environmental impact? Trends Parasitol 26:44–51.

Watt KA, Nussey DH, Maclellan R, Pilkington JG, McNeilly TN. 2016. Fecal antibody levels as a noninvasive method for measuring immunity to gastrointestinal nematodes in ecological studies. Ecol Evol 6:56–67.

Wilson K, Grenfell BT, Pilkington JG, Boyd HEG, Gulland FMD. 2004. Parasites and their impact. In: T. Clutton-Brock and J. Pemberton, editor. Soay Sheep: Dynamics and Selection in an Island Population Cambridge University Press. p. 113–65.

Wood CL, Lafferty KD. 2013. Biodiversity and disease: a synthesis of ecological perspectives on Lyme disease transmission. Trends Ecol Evol 28:239–47.

Zuur AF, Ieno EN, Saveliev AA. 2017. Beginner’s guide to spatial, temporal, and spatial-temporal ecological data analysis with R-INLA.

